# Buffer tolerance landscape of LbCas12a trans-cleavage efficiency

**DOI:** 10.64898/2026.09.23.753913

**Authors:** Roberto Alcántara, Roberto Spurio, Pohl Milón

## Abstract

Numerous CRISPR-Cas systems have been developed for molecular detection of genetic elements exploiting the trans-cleavage activity of LbCas12a combined with short fluorescent probes. Alongside, a large variability of buffer conditions has been reported. However, how solution chemistry balances enzymatic turnover, signal stability, and detection performance have not been systematically defined. Here, we evaluated how the interplay between anions (chloride *vs*. acetate), additives, sequence probes, and ionic strength contributes to LbCas12a trans-cleavage activity. Buffer composition screening revealed that chloride-based buffers maintain a low background but slow catalytic rates, meanwhile acetate-based buffers accelerate enzymatic turnover with and without the target DNA sequence, increasing non-specific signals. Reducing agents and surfactants were the best enhancer combinations, improving cleavage performance by increasing ternary complex formation (*K*_m_). High salt concentrations only slightly improved the reaction kinetics. Finally, the 5’FAM-TTATT-3’BHQ-1 reporter showed the best balance between signal intensity and nonspecific background. Our findings indicate that LbCas12a tolerates a vast buffer composition landscape, maintaining robust trans-cleavage activity and detection performance. However, conditions that enhance catalytic activity simultaneously increase nonspecific probe cleavage and higher background, resulting in lower detection performance. Therefore, assay optimization should prioritize detection performance rather than maximal fluorescence output, where solution chemistry is a critical determinant.

## INTRODUCTION

CRISPR-Cas systems include diverse protein-RNA complexes with nuclease activity, found across many bacterial and archaeal genera^1–3^. Originally discovered as an adaptive immune mechanism against exogenous nucleic acids such as plasmids and phages, these systems have become widely used in biotechnology due to their relatively simple target recognition mechanism^4,5^. Molecular recognition is achieved through base-pair hybridization between the target (i.e., DNA or RNA) and the crRNA, which guides the CRISPR-Cas complex to its complementary sequence^6–8^. Upon target recognition, a series of conformational rearrangements enable the target and non-target strands to access LbCas12a catalytic site, where cleavage occurs^9–11^. Some CRISPR-Cas complexes, especially those belonging to Class II (Types II, V and VI), also exhibit nonspecific cleavage of nearby DNA or RNA molecules once activated^7,12,13^. Although no *in vivo* advantages have been clearly established for this trans-cleavage activity, it has been extensively exploited for nucleic acid detection platforms^10,14–19^.

To date, all CRISPR-based molecular detection assays rely on this collateral trans-cleavage activity, using nucleases such as Cas12, Cas13 and more recently Cas14^16,17,20,21^. In these assays, a reporter molecule (DNA or RNA) is labeled with a signal-generating pair, which may consist of a fluorophore and quencher, a fluorophore and biotin, a fluorophore and reactive groups, or electrochemical tags^22–24^. When an activated CRISPR-Cas complex cleaves the reporter, a detectable signal is produced that strictly correlates with the presence and quantity of the target^10,25–28^. As for other detection platforms based on direct or indirect signal generation, a primary strategy for improving assay performance is to enhance the efficiency of the signal-producing mechanism. Consequently, numerous studies have explored Cas and crRNA engineering, probe design, electrochemical- and nanomaterial-based readouts, and reaction buffer optimization^28–32^.

Buffer optimization is certainly a critical step of any assay involving enzymatic reactions. For CRISPR-Cas-based assays, there exists a myriad of buffer formulations and reporter probes^33,34^. This, in turn, brings a high level of complexity for comparing and reproducing detection boundaries among different reports. Although some studies have compared commonly used buffer or benchmarked conditions from earlier reports^35–38^, a systematic, stepwise evaluation of how individual buffer components influence Cas trans-cleavage activity is still lacking. This gap limits our understanding of how specific chemical factors shape the overall reaction landscape, sensitivity, and specificity.

In this study, we identified the most frequently reported buffer components across published CRISPR-Cas assays and performed a systematic characterization of the interplay between buffers, additives, ionic strength and probes on the trans-cleavage activity of LbCas12a ortholog using a short dsDNA target. We aimed to determine which buffer components influence trans-cleavage activity as observed by raw fluorescence and signal-to-background ratios (indicative of molecular specificity) and therefore which ones should be prioritized when designing or optimizing CRISPR-Cas detection assays.

## RESULTS

### Trans-cleavage kinetics of the LbCas12a depend on buffer composition

The unspecific trans-cleavage activity is a key feature for using Cas12, Cas13 and, recently, Cas14 nucleases in the design of next-generation molecular detection assays. However, the kinetic profile of this reaction is heavily affected and modulated by the solution chemistry. To investigate the effect of buffer composition on the trans-cleavage activity, we first evaluated the impact of the fundamental chemical environment on reaction kinetics and detection performance, focusing on raw fluorescence, detection response (NF_ntc_, defined as the fluorescence ratio relative to the non-template control (NTC) reaction), and initial velocity (V_0_). Univariate analysis revealed a key difference between the two most common buffers, at the same pH = 7.9. The Tris-Ac^-^ buffer promoted a faster enzymatic activity, reaching raw fluorescence signal saturation between 30 and 40 minutes, on average, for all additives (Figure 1A, Figures S2-3). Despite this, a time-dependent increase in the raw fluorescence of the NTC reaction was observed typically after 30 minutes under all tested conditions (Figure 1A, Figure S2-3). Nevertheless, because typical assay readout times range from 5 to 30 minutes, this late increase in the unspecific fluorescence is expected to have minimal impact on diagnostic interpretation. In contrast, the Tris-Cl^-^ buffer showed a slower reaction with a less pronounced fluorescence signal in the NTC reaction signal, yielding a more stable kinetic profile throughout a 90-minute reaction (Figure 1A, Figure S2-3).

**Figure 1.**
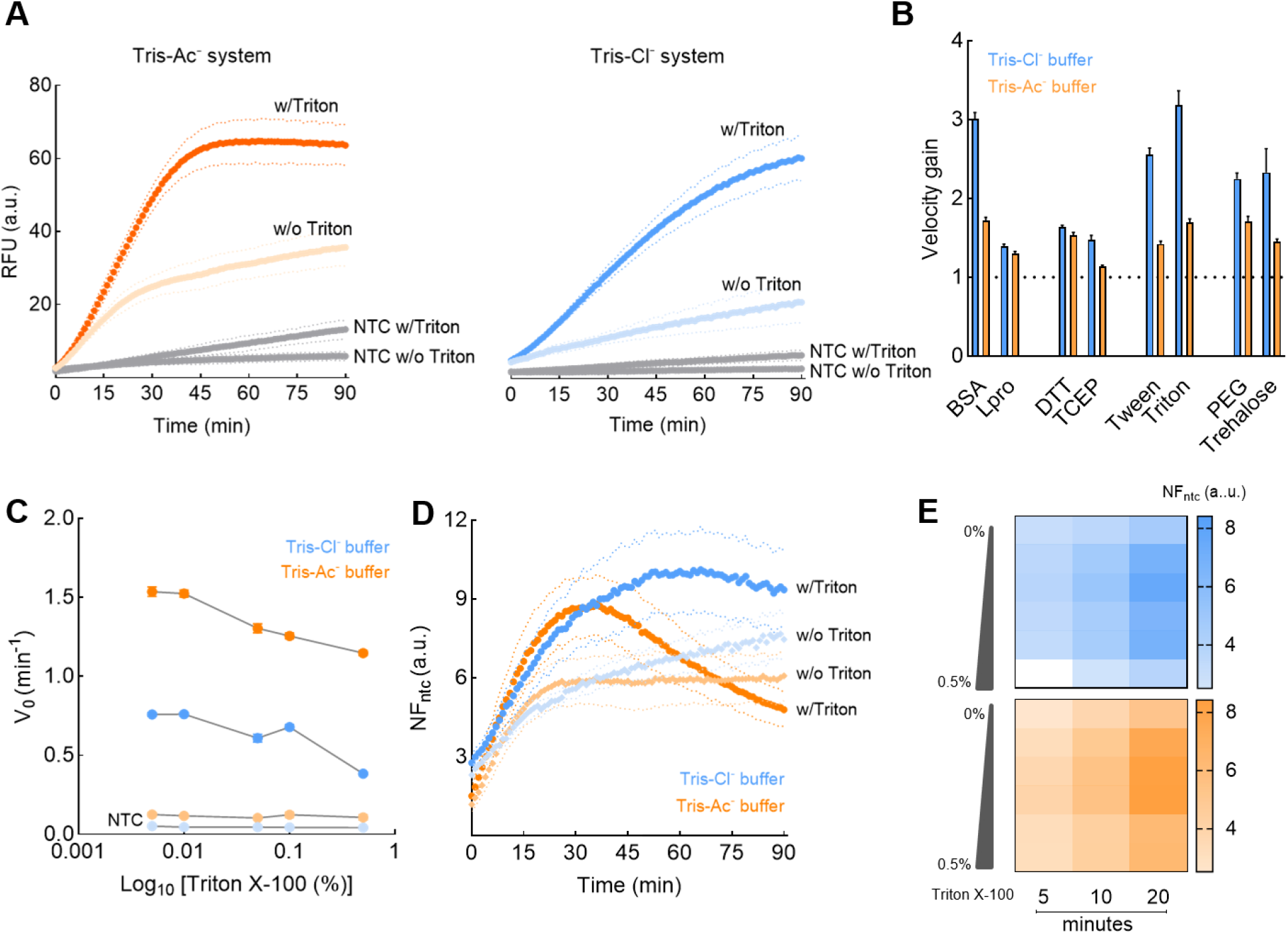
Univariate screening of different additive groups for the trans-cleavage activity of LbCas12a. Additives were grouped based on the expected effect on enzyme activity such protein stabilizer (BSA, L-proline), reducing agents (DTT, TCEP), surfactants (Tween 20, Triton X-100), and crowding agents (PEG-8000, trehalose). The effect of each additive was evaluated in two different buffers: Tris-Cl^-^ (10 mM Tris-HCl, 50 mM NaCl; blue) and Tris-Ac^-^ (20 mM Tris-AC, 50 mM CH_3_CO_2_K; orange). (A) Representative fluorescence time courses of LbCas12a with 0.005% Triton X-100 compared to non-additive and non-template controls (NTC). Dotted lines represent the standard deviation (n=3). (B) Velocity gain for LbCas12a reactions in buffer supplemented with the additives at their lowest concentrations. Data is normalized to the non-additive control (Y = 1, dashed line) for Tris-Cl- and Tri-Ac-buffers. (C) Dose-response analysis of Triton X-100. The reaction exhibits an inhibition response, where concentrations exceeding 0.01% decrease the initial velocity (V_0_) in both buffers. (D) Same as A, but for detection response (i.e., NF_ntc_, fluorescence ratio relative to the NTC control). Tris-Ac-buffer yields a rapid increase followed by a decrease, whereas Tris-Cl^-^ maintains signal stability over 90 minutes. Dotted lines represent the standard deviation (n=3). (E) Heatmap summary of detection response values at defined readout times (5, 10, and 20 min) across the Triton X-100 concentration gradient (0 to 0.5%), in Tris-Ac^-^ buffer (orange) versus Tris-Cl^-^ (blue), showing the higher signal increase in Tris-Ac^-^ buffer in a typical detection window.

To study the effect on the trans-cleavage kinetics of chemical enhancers, they were grouped by their functional role: protein stabilizers (BSA, L-proline), reducing agents (DTT, TCEP), surfactants (Triton X-100, Tween 20), and crowding agents (PEG8000, trehalose). At the lowest concentration tested, surfactants such as Triton X-100 and protein stabilizers such as BSA produced the strongest enhancement of trans-cleavage activity. In the Tris-Cl^-^ buffer, these compounds increased the initial velocity (V_0_) by approximately 3-fold compared to the non-additive control (Figure 1B). While enhancements were also observed in the Tris-Ac^-^ buffer, the relative fold-change was less pronounced. Dose-response analysis showed that kinetic enhancement is strictly concentration-dependent. For Triton, higher concentrations showed an inhibitory effect on the initial velocity (Figure 1C). With the exception of L-proline and trehalose, the remaining additives showed a marked inhibitory effect (Figure S4). In all cases, Tris-Ac-consistently produced higher fluorescence signals than Tris-C^l-^.

Analysis of the detection response showed that reactions in the Tris-Ac^-^ buffer reach a peak early, but then declines over time as the unspecific fluorescence signal from the NTC reaction rises. This effect is clearly observed for BSA, Triton X-100, Tween20 and PEG800 (Figure 1D, Figures S2-3). Conversely, Tris-Cl^-^ conditions maintained a lower but stable kinetic profile. Evaluating the detection response within a typical window for rapid end-point assays (5–20 min) evidenced that Tris-Ac^-^ buffer offers slightly higher detection resonse compared to Tris-Cl^-^ (Figure 1E). All together, these results demonstrate that LbCas12a tolerates a wide range of additives in buffer compositions while retaining trans-cleavage activity.

### Reducing agents and surfactants stand out as synergistic enhancers

Based on univariate analysis results, we explore how combinations of top-performing candidates (BSA, DTT, Triton X-100, and PEG-8000) could induce synergistic enhancement. A consistent kinetic profile was observed if compared to the univariate analysis. Tris-Ac^-^ buffer showed a higher raw fluorescence reaching plateau at 40 min on average, for all conditions. The rise of the unspecific fluorescence was more intense in the presence of any chemical enhancer combination, for either reducing or crowding agents, after 20 min on average. In contrast, the Tris-Cl^-^ buffer maintained baseline stability but failed to match the catalytic velocity of the Tris-Ac^-^ buffer (Figure 2A, Figure S5). Analysis of initial velocity (V_0_) showed that combinations including a reducing agent and a surfactant increased the initial velocity compared to the standard NEB r2.1 buffer (Figure 2B). These combinations, such as DTT:PEG and DTT:Triton:PEG, outperformed the others in both buffers (Figure 2B). End-point analysis at 20 min revealed that high detection response was obtained in combinations including either DTT, Triton X-100 or PEG 8000 (Figure 2C).

**Figure 2.**
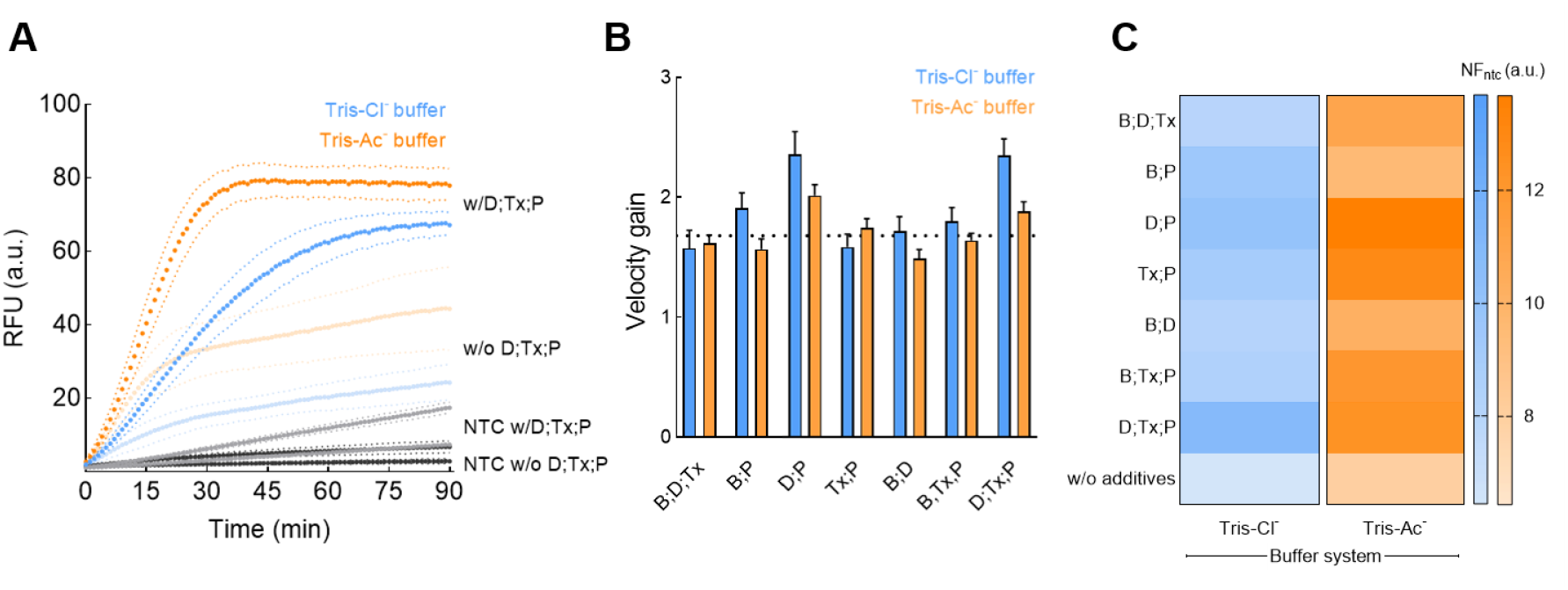
Combinatorial screening of buffer additives for synergistic enhancement on the LbCas12a trans-cleavage activity. Based on univariate analysis, top-performing additives BSA (B), DTT (D), Triton X-100 (Tx), and PEG 8000 (P) were selected. (A) Raw fluorescence kinetics of the top-performing combination (DTT:Triton:PEG – D;TX;P), compared to non-additive controls. Dotted lines represent the standard deviation (n=3). (B) Normalized initial velocity (V_0_) of LbCas12a reactions. Initial velocity was estimated in the linear range (< 20 min) and normalized against the non-additive control. The dotted line in the Y-axis marks the enhancement factor on the trans-cleavage activity for the LbCas12a in the standard buffer NEB r2.1 (100 µg/mL BSA). Two conditions showed an improvement on the initial velocity: DTT:PEG (D;P) and DTT:Triton:PEG (D;Tx;P), in both buffers, Tris-Cl^-^ and Tris-Ac^-^. (C) Heatmap of detection response (NF_ntc_) at 20 min of reaction.

Comparison between both buffers showed that DTT:Triton:PEG and DTT:PEG produced the highest detection response. To resolve this interplay, we performed a checkerboard titration in the Tris-Ac^-^ buffer. Evaluation of the DTT:Triton ratio revealed a narrow optimal window at lower enhancer concentration with DTT at 1.0 mM and Triton X-100 at 0.005% as the best combination (Figure S6). As expected, higher concentrations of PEG 8000 showed an antagonistic effect, resulting in a linear inhibition of the initial velocity (V_0_). Minimal difference in detection response was observed with PEG supplementation (Figure S6). Consequently, the crowding agent was excluded for downstream characterization.

### Reporter sequence did not modulate the trans-cleavage performance

Beyond buffer chemistry, the sequence and topology of the ssDNA reporter have been considered as determinants of the trans-cleavage efficiency. Therefore, we evaluated a panel of probes, tested in the supplemented Tris-Ac^-^ buffer, to study how probe engineering affects trans-cleavage activity. Raw fluorescence analysis showed that higher fluorescence intensity was obtained with the C-rich modified (TTATT5C) and the poly-T (15T) probes. However, these increased signals co-occur with an amplified unspecific fluorescence signal in the NTC controls, ranging from 2-fold for the poly-T probe to 4-fold for the modified standard probe compared to the canonical TTATT probe (Figure S7). Detection response analysis confirmed that TTATT probe yielded the best performance, with values ranging from a 1.5-fold increase over the poly-T (15T) probe to a 3-fold increase compared with the hairpin probe at 20 min of fluorescence measurement. All probes, with exception of the hairpin, peaked within 20 min of reaction (Figure S7). The presence of cytosine residues (8C and TTATT5C) slowed down the reaction compared to T-rich probes. Collectively, these results indicate that longer probes may produce higher apparent trans-cleavage yield, yet also the non-specific reaction (NTC) is enhanced, resulting in lower detection response values, therefore loss of detection performance. Consequently, the short unstructured probe was retained for subsequent analysis

### Optimized Tris-acetate buffers accelerate target detection and improve analytical sensitivity

The effect of ion concentration on the trans-cleavage activity was evaluated by a checkerboard titration in the additive-supplemented buffer. The screening revealed a positive tendency between ionic strength and the raw fluorescence signal. Reactions in 75 mM CH_3_CO_2_K showed on average the best conditions for trans-cleavage performance, with the highest detection response observed at 25 mM Mg(CH_3_CO_2_)_2_ (Figure 3A). Comparison with the previous ion conditions (50 mM CH_3_CO_2_K and 20 mM Mg(CH_3_CO_2_)_2_showed that while the raw fluorescence signal remained comparable during the initial linear phase, the higher ionic strength flattened the unspecific fluorescence slope in the late reaction phase (Figure 3B) which resulted in a 20% increase in the normalized signal output.

**Figure 3.**
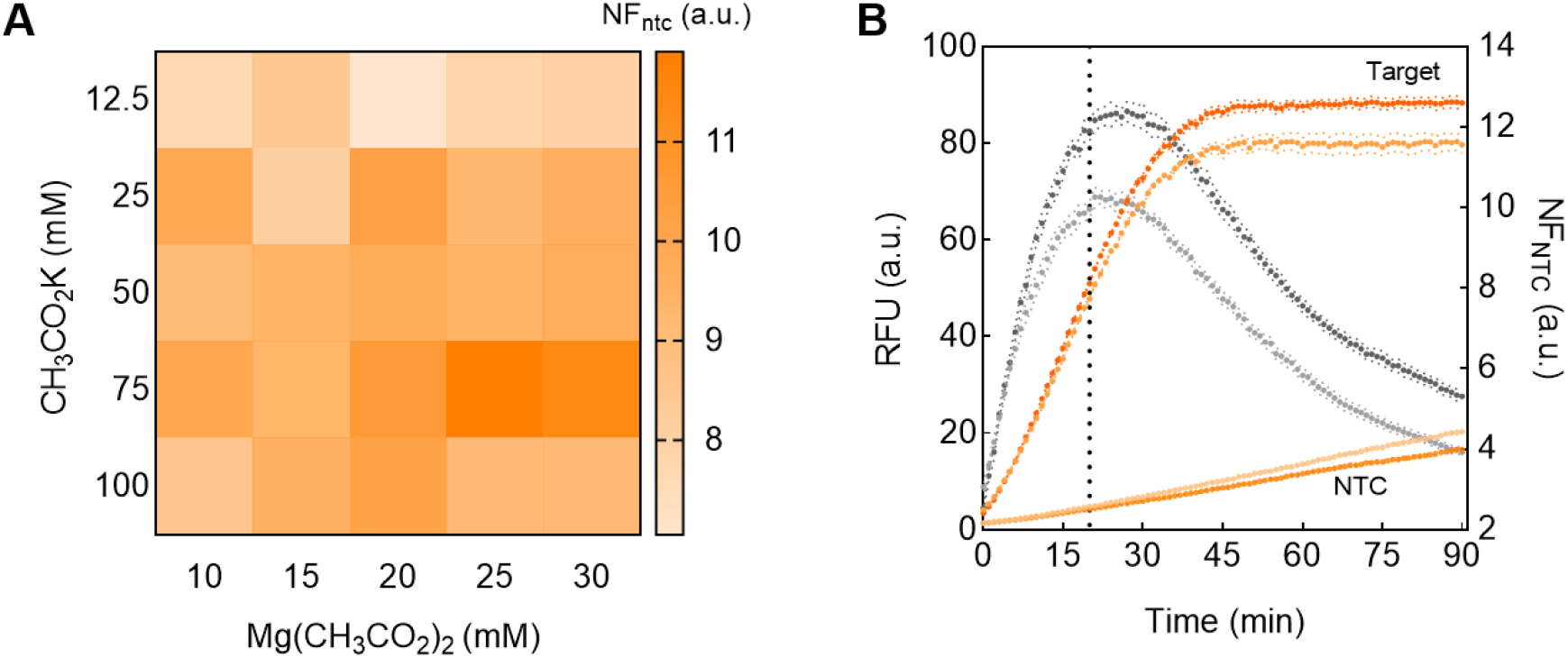
Ionic strength effect on the LbCas12a trans-cleavage activity on a DTT-Triton-supplemented Tris-Ac^-^ buffer. (A) Checkerboard screening of magnesium acetate (Mg(CH_3_CO_2_)_2_, 10-30 mM) and potassium acetate (CH_3_CO_2_K, 12.5-100 mM) in the optimized buffer containing 1.0 mM DTT and 0.005% Triton X-100. Heatmap intensity represents the detection response values (NF_ntc_) at 20 minutes. (B) Comparative kinetic profiles of the previous tested Tris-Ac^-^ condition (50 mM CH_3_CO_2_K with 20 mM Mg(CH_3_CO_2_)_2_; light dotted lines) versus the identified best high-salt condition (75 mM CH_3_CO_2_K with 25 mM Mg(CH_3_CO_2_)_2_; dark dotted lines). The relative fluorescence units (RFU) correspond to the orange curves for the positive reaction against the non-template control (NTC). Meanwhile the detection response values (NF_ntc_) are presented as the grey curves. Before 20 minutes (dotted line in the X-axis), no differences in RFU were observed. However, the higher ionic strength increased by 20% the observed NF_ntc_ values.

To evaluate how these integrated factors affect analytical performance, we compared them with the standard Tris-Cl^-^ buffer. Detection response analysis across a target titration gradient revealed distinct kinetic profiles. Reactions in the Tris-Ac^-^ buffer (both additive-only and salt-optimized) showed a faster kinetic, peaking within a 20-minute window, compared with the Tris-Cl^-^buffer, which peaked at ∼30 min (Figure 4A-C). A two-fold increase was observed at the lowest detectable target concentration, decreasing from 78 pM in the Tris-Cl^-^ buffer to 19 pM in the high-salt Tris-Ac^-^ buffer. Optimized Tris-Ac^-^ buffers also showed lower Michaelis constants (*K*m = 0.28 – 0.38 nM) compared with the Tris-Cl^-^ buffer (*K*m = 1.14 nM) (Figure 4D). The limit of detection (LOD) ranged from 19 pM for the additive-supplemented Tris-Ac^-^ buffer to 36 pM for the Tris-Cl^-^ buffer, respectively. The use of high salt Tris-Ac^-^ buffer showed a LOD below 9 pM, although it could not be precisely estimated under the current results (Figure S8). Importantly, unspecific fluorescence signal was not particularly affected by any of the tested buffer formulations in non-cognate and non-template reactions, confirming that the observed effects were specific to the presence of the cognate target (Figure S9).

**Figure 4.**
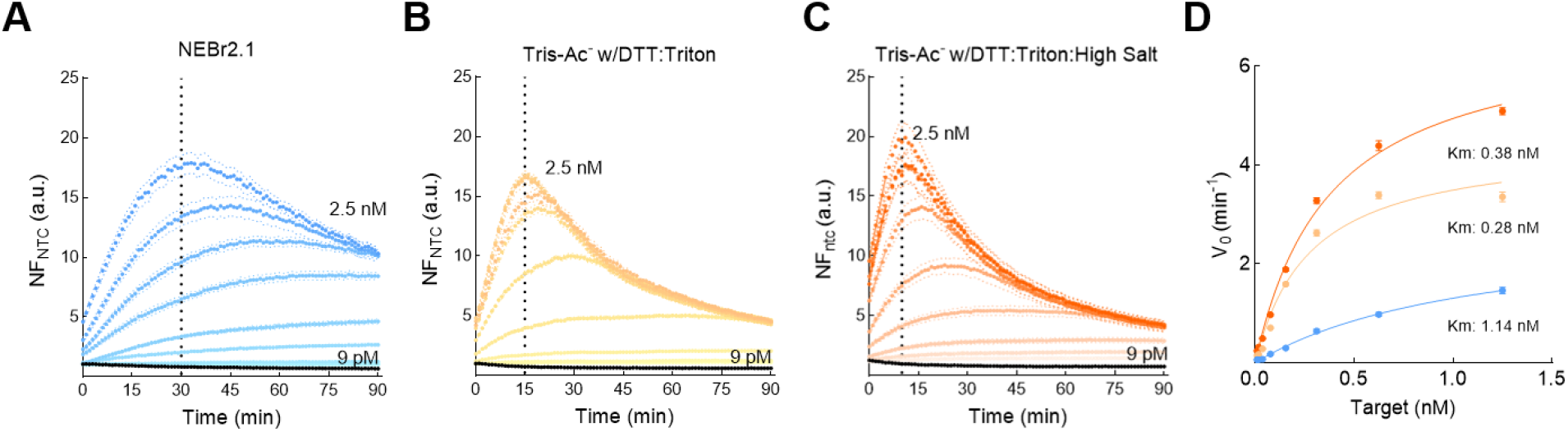
Analytical sensitivity and kinetic efficiency of an optimized buffer for LbCas12a trans-cleavage activity. (A–C) Detection response (NF_ntc_) over time across a target titration gradient (0.10 – 1.25 nM). Reactions were performed in: (A) standard NEB r2.1 buffer, (B) Tris-Ac^-^ buffer (+DTT/Triton), and (C) optimized high-salt Tris-Ac^-^ buffer (+DTT/Triton/High salt). The black dotted line marks the time when the detection response reaches its peak value, for Tris-Cl^-^ buffer at 30 min, for Tris-Ac^-^ buffer at 15 min, and for the optimized high-salt Tris-Ac^-^ at 10 min. At the 20-min readout, analytical sensitivity improved in a stepwise manner, with the lowest detectable target concentration decreasing approximately two-fold between successive buffers: high-salt Tris-Ac− < low-salt Tris-Ac− < NEBuffer r2.1. Black lines at the bottom of each graph represents the NF_ntc_ values reached with maximum concentration of a non-cognate template. (D) Michaelis-Menten kinetic analysis of LbCas12a trans-cleavage rates as a function of target quantity. The final optimized high-salt Tris-Ac^-^ buffer (dark orange) exhibits a higher binding affinity (*K*_m_ = 0.38 nM), considerably outperforming the standard Tris-Cl^-^ buffer (blue, *K*_m_ = 1.14 nM). Error bars represent standard deviation (n=3).

Taken together, this data demonstrates that using Tris-Ac^-^ buffer result in higher raw fluorescence outputs within a typical detection window, aligned with most publications where CRISPR-Cas complexes are used as molecular detection systems. Additionally, the chemical background assumes a critical role in determining the efficiency of the trans-cleavage activity outweighing the impact of reporter probe engineering. These findings suggest the CRISPR-Cas detection reaction is tolerant to a wide range of buffer compositions; and simple, rationally designed combinations are sufficient to maximize the fluorescence output; however, detection performance can greatly vary particularly with reaction time.

## DISCUSSION

Molecular diagnostics based on CRISPR-Cas systems have emerged as a highly adaptable platform for accurate, specific, affordable, and field-deployable molecular detection^39–41^ of pathogens and disease-related genetic markers^24,42–45^. However, there is a wide range of optimal reaction conditions described^33– 38^, making performance comparison difficult, which in turn challenges the translational aspects of this technology, from laboratories to point-of-care settings. To address this gap, this study systematically evaluated the effect of common anions, chemical additives, reporter probe structures, and ionic strength on the LbCas12a trans-cleavage activity.

Across all chemical environments evaluated, the LbCas12a consistently exhibited trans-cleavage activity, generating robust but variable fluorescence signals. This broad functional tolerance confirms that no single chemical environment is exclusively optimal for CRISPR-Cas12a-based detection. Baseline anion composition was identified as the primary determinant of raw fluorescence output and catalytic velocity. Tris-Ac^-^ buffers consistently produced the highest fluorescence and fastest initial velocity (V_0_), even in the absence of chemical additives. These findings empirically validate protocols for nucleic acid detection favoring Tris-Ac^-^ reaction buffers^35,37,46,47^.

However, Tris-Ac^-^ buffers induced a progressive rise in nonspecific background fluorescence (non-template reactions) markedly after 20 minutes, likely due to acetate-mediated alterations in the hydration shell of the enzyme or thermodynamic flexibility that induce a destabilization of the inactive Cas:crRNA complex, exposing the RuvC catalytic domain and enabling target-independent probe degradation. This kinetic dichotomy represents a risk for false-positive results if readout time windows are not strictly managed, particularly in low-target samples.

Screening of chemical additives showed that non-ionic surfactants and reducing agents synergistically enhanced fluorescence output. Sequence analysis shows that LbCas12a contains nine conserved cysteine residues, supporting the activating effect of reducing agents, which likely maintain solvent-exposed sulfhydryl groups essential for structural stability^46,48^. Non-ionic surfactants such as Triton X-100 exert non-specific effects by preventing enzyme agregation and/or adsorption to hydrophobic surfaces, in that way maintaining higher concentrations of active complexes in solution. These results align with previous reports^32,36,37,49,50^, demonstrating improved Cas12 activity upon inclusion of those reagents.

Crowding agents like PEG-8000 also enhanced activity, presumably by stabilizing in-solution complexes^35,49^. However, their combination with DTT and Triton X-100 did not further increase signal output, suggesting that simple additive formulations are sufficient to increase the fluorescence signal. Very low concentrations of additives (e.g., 1.0 mM dithiothreitol (DTT) and 0.005% Triton X-100), were found optimal to achieve best enzymatic performance, whereas higher concentrations caused progressive linear inhibition.

Ionic strength also modulates enzymatic activity by stabilizing protein structure and influencing substrate binding through the interplay of electrostatic interactions^51,52^. Divalent Mg^2+^ ions, essential cofactors for endonuclease activity^32,53–55^, positively correlated with fluorescence output, consistent with prior studies showing enhanced Cas12a activity above the standard 10 mM concentration^46,49^. Monovalent ions (Na^+^, K^+^) have been hypothesized to influence PAM sequence recognition and substrate access to the RuvC pocket. The positively charged α-helical lid of the NUC domain facilitates interaction of the negatively charged backbone of target and reporter probes into the RuvC catalytic pocket^32,37,56,57^. At high concentrations, cations may hamper probes access to the catalytic pocket. This effect has mainly observed in Tris-Cl^-^ buffers, and appears to be ortholog-dependent, with LbCas12a being comparatively less sensitive ^32,57^. Our results showed a positive correlation between fluorescence and K^+^ concentration, aligned with previous results where optimal monovalent ion concentration is at least 50 mM^37,46,53,58^. Although the role of monovalent ions remains controversial, high ionic strength may provide electrostatic shielding that suppresses nonspecific low-affinity, non-specific probe cleavage^57^.

Reporter probe design is another critical factor. Because probes must access the catalytic site while the ternary CRISPR:Cas:target complex remains assembled, sequence and structure can influence RuvC interactions^8,11,59–61^. Previous studies reported enhanced performance for C-rich extensions (TTATT5C)^62^ or homopolymeric probes (8C and 15T)^37,46,58,63^. However, under our experimental conditions, these probes simultaneously generated up to 4-fold increase in nonspecific signal compared to the standard short probe (TTATT). Structured probes, such as the 10T-hairpin, did not yield improved fluorescence output as reported previously^37,64^. Notably, performance differences described previously were obtained in different buffers, most based on Tris-Cl^-^ buffers. In contrast, our results show that the simple, unstructured TTATT reporter achieved the highest overall signal-to-blank ratio (NF_ntc_), highlighting that rational buffer evaluation provides better outcomes compared to complex probe engineering.

### Limitations of the study

This work focused exclusively on the LbCas12a ortholog, and the findings may not be directly generalizable to other Cas12 variants with distinct biochemical properties. Although buffer compositions and additives were systematically screened based on a literature-guided selection, the high tolerance observed suggests that untested reagents are unlikely to drastically alter catalytic activity. Finally, short linear double-stranded DNA targets were employed to simulate nucleic-acid-amplification assay outcomes, but these substrates may not fully capture the structural complexity of plasmid or genomic DNA in amplification-free detection formats.

## RESOURCE AVAILABILITY

### Lead contact

Requests for further information and resources should be directed to and will be fulfilled by the lead contact, Pohl Milon.

### Materials availability

This study did not generate new unique reagents.

### Data and code availability

- All data reported in this paper will be shared by the lead contact upon request.
- This paper does not report original code.
- Any additional information required to reanalyze the data reported in this paper is available from the lead contact upon request.

## ACKNOWLEDGMENTS

This study was supported by the Universidad Peruana de Ciencias Aplicadas with grant C-005-2023 to RA.

## AUTHOR CONTRIBUTIONS

Conceptualization, R.A. and P.M.; methodology, R.A.; Investigation, R.A.; writing – original draft, R.A. and P.M.; writing – review & editing, R.A., R.S. and P.M.; funding acquisition, R.A.; resources, P.M.; supervision, R.S., and P.M.

## DECLARATION OF GENERATIVE AI AND AI-ASSISTED TECHNOLOGIES IN THE WRITING PROCESS

During the preparation of this work, the author(s) used Gemini (AI assistant) to improve grammar, readability, and clarity of the manuscript. After using this tool, the author(s) carefully reviewed and edited the content as needed and take full responsibility for the accuracy and integrity of the publication.

## SUPPLEMENTAL INFORMATION

Document S1. Supplemental Figures S1–S9

## STAR★METHODS

### KEY RESOURCES TABLE

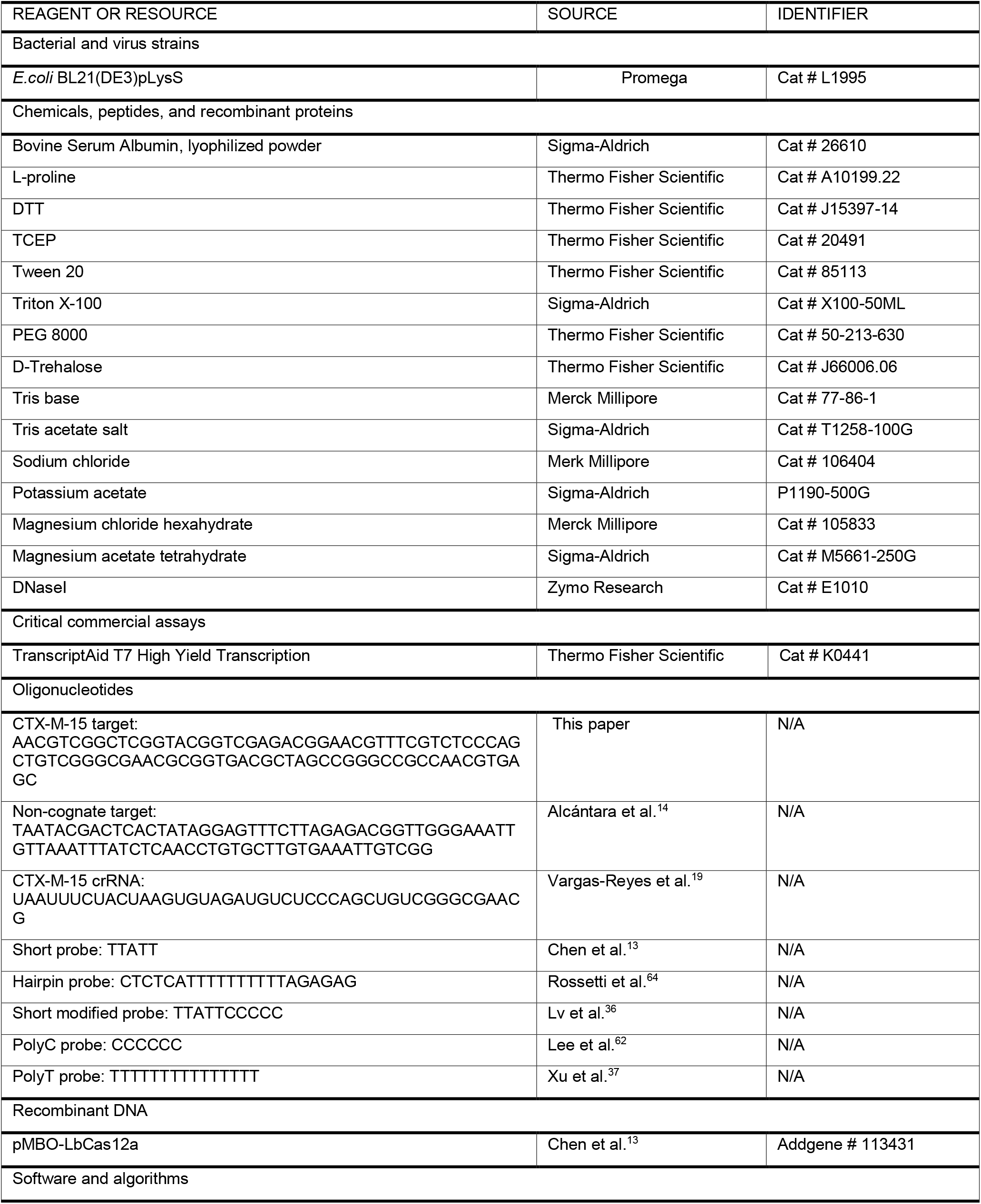
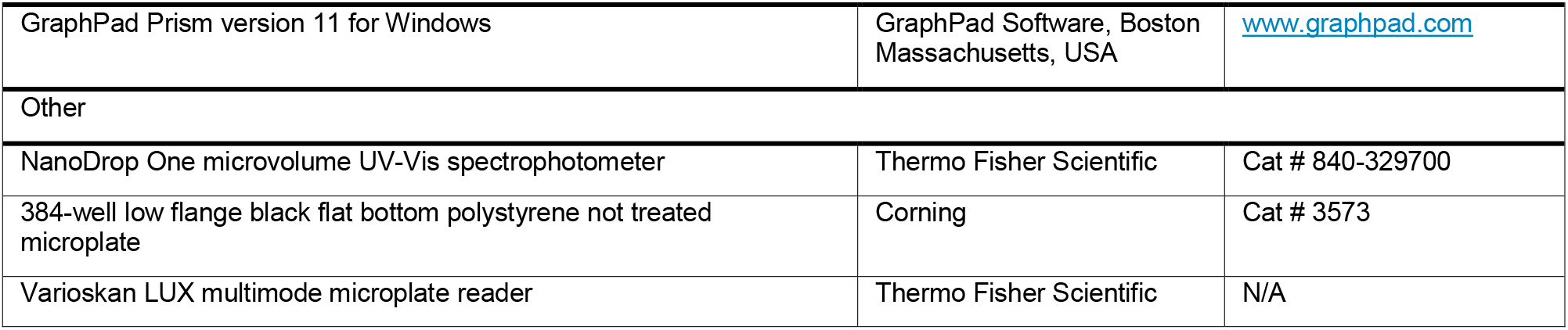

## METHOD DETAILS

Based on a literature analysis, reaction conditions potentially affecting LbCas12a trans-cleavage activity were systematically evaluated through a sequential screening workflow (Figure S1). All assays were performed in triplicate unless otherwise indicated.

### Biological stocks

LbCas12a, derived from *Lachnospiraceae bacterium*^*13*^ was produced from a cryopreserved *E. coli* BL21(DE3)pLysS harboring the pMBP-LbCas12a plasmid, following our previously published protocol^65^. For all CRISPR-Cas assays, we used our previously reported crRNA specific for *blaCTX-M-15* gene^19^. The crRNA was produced by *in vitro* transcription using the TranscriptAid T7 High Yield Transcription Kit. A double-stranded DNA including the promoter T7 and the crRNA sequence was used as a template for transcription. Double-labeled probes carried a 5’FAM fluorophore and 3’BHQ1 quencher. As the target sequence, we used a 89-bp double-stranded DNA fragment containing the *blaCTX-M-15* region of interest. Additionally, a non-cognate target without the target sequence yet using the same PAM sequence (TTTC) as the cognate target was included as a specificity control. All probes and target DNA fragments were purchased from Macrogen Inc. (Seoul, South Korea).

### CRISPR-Cas reactions

CRISPR-Cas reactions were prepared as previously reported^14^. Typically, 1 µM crRNA was activated by heating at 65°C for 10 minutes in nuclease-free water and then allowed to refold at room temperature for 10 min. Next, 50 nM LbCas12a was mixed with 75 nM of the refolded crRNA in the corresponding buffer (Tris-Cl^-^ at pH 7.9 or Tris-Ac^-^ at pH 7.9). As control for performance comparison, the standard NEBr2.1 (10 mM Tri-Cl^-^, pH 7.9; 50 mM NaCl, 100 µg/mL BSA) buffer was used. The labeled probed (1 µM) was added to the complex, and the mixture was incubated in dark for 10 minutes. The dsDNA target (200 pM) (i.e., cognate or non-cognate sequence) was prepared in the corresponding buffer with supplemented Mg^2+^ (i.e., MgCl^2^ or Mg(CH^3^CO^2^)^2^, as appropriate). Finally, a 50 µL reaction was prepared consisting of 10 nM LbCas12a, 15 nM crRNA and 200 pM dsDNA target. A non-template control was included in all assays to calculate the normalized fluorescence signal in the absence of any analyte (NF_NTC_). Reactions were prepared in a 384-well low flange, black, flat-bottom microplate (Corning, cat. N° 3573) and read on a Varisokan Lux plate reader using the following settings: excitation at 491 nm, emission at 525 nm, with fluorescence measurements taken every 1 min for 90 min.

Fluorescence output was analyzed both as raw fluorescence and as normalized fluorescence to the non-template control. Initial velocity (V_0_) was calculated using linear regression from the linear region of each time course for all experimental conditions.

### Evaluation of the role of additives on trans-cleavage activity

The additives evaluated included protein stabilizers (e.g., BSA and L-proline), reducing agents (e.g., DTT and TCEP), surfactants (e.g., Tween 20 and Triton X-100), and crowding agents (e.g., PEG 8000 and trehalose). Each additive was first evaluated independently in both buffers. For each additive, a concentration curve was prepared relative to the non-additive control. Additives were added to the corresponding buffer used to dilute the dsDNA target and the mixture was then mixed with the Cas:crRNA:probe complex as indicated above. The tested concentration ranges were as follow: 50 – 250 µg/mL BSA, 0.1 – 0.5 M L-proline, 0.5 - 2.5 mM DTT, 0.1 - 1 mM TCEP, 0.01 - 1% Tween 20, 0.005 - 0.5% Triton X-100, 0.1 - 4% PEG 8000, and 1 - 5% D-trehalose.

Based on the additive screening results, the concentration of one additive from each group that yielded the highest fluorescence signal (i.e., initial velocity and normalized fluorescence) was selected, including 50 µg/mL BSA, 1.5 mM DTT, 0.005% Triton X-100, and 0.1% PEG-8000. Additive mixtures were then prepared to include two- and three-component combinations, along with a non-additive control. All mixtures were evaluated in both buffers. For the mixtures that exhibited the best performance, a checkerboard assay was subsequently performed. In this assay, all additives present in the top-performing mixtures, regardless of whether they originated from different combinations, were evaluated across the full concentration ranges described in the previous section.

Once the optimal additive mixture was identified, a checkerboard assay was performed to evaluate the effects of monovalent and divalent ions on the Tris-Ac^-^ buffer. The monovalent ion (K^+^) was tested at concentrations ranging from 12.5 – 100 mM, while the divalent ion (Mg^2+^) was evaluated from 10 – 30 mM.

### Analytical sensitivity

Signal dependence on target concentration was evaluated using DNA concentrations ranging from 0.01 – 2.5 nM in 50 µL reactions. The CRISPR-Cas assay was performed under three buffer conditions: (i) NEBuffer r2.1 (pH 7.9), and in two Tris-Ac^-^ buffers (pH 7.9) supplemented with DTT-Triton X-100, containing either (ii) 20 mM Mg(CH_3_CO_2_)_2_ – 50 mM CH_3_CO_2_K, or (iii) 25 mM Mg(CH_3_CO_2_)_2_ – 75 mM CH_3_CO_2_K

## QUANTIFICATION AND STATISTICAL ANALYSIS

Relative fluorescence units were analyzed both as raw fluorescence units (a.u.), and as normalized fluorescence ratios (NFntc), calculated as the ratio between the raw fluorescence of the positive reaction and that of the non-template control (NTC). Initial velocity was estimated as the slope obtained by nonlinear regression of the linear region of the fluorescence curve from 0 to 20 minutes. Kinetic parameters (*K*m and Vmax) were calculated by non-linear regression using the Michaelis-Menten equation. All analysis and graphics were performed using GraphPad Prism, version 10.6.0. Standard deviation and standard error of the mean were calculated when required.

