## Supplementary figures for "Buffer tolerance landscape of LbCas12a trans-cleavage efficiency"

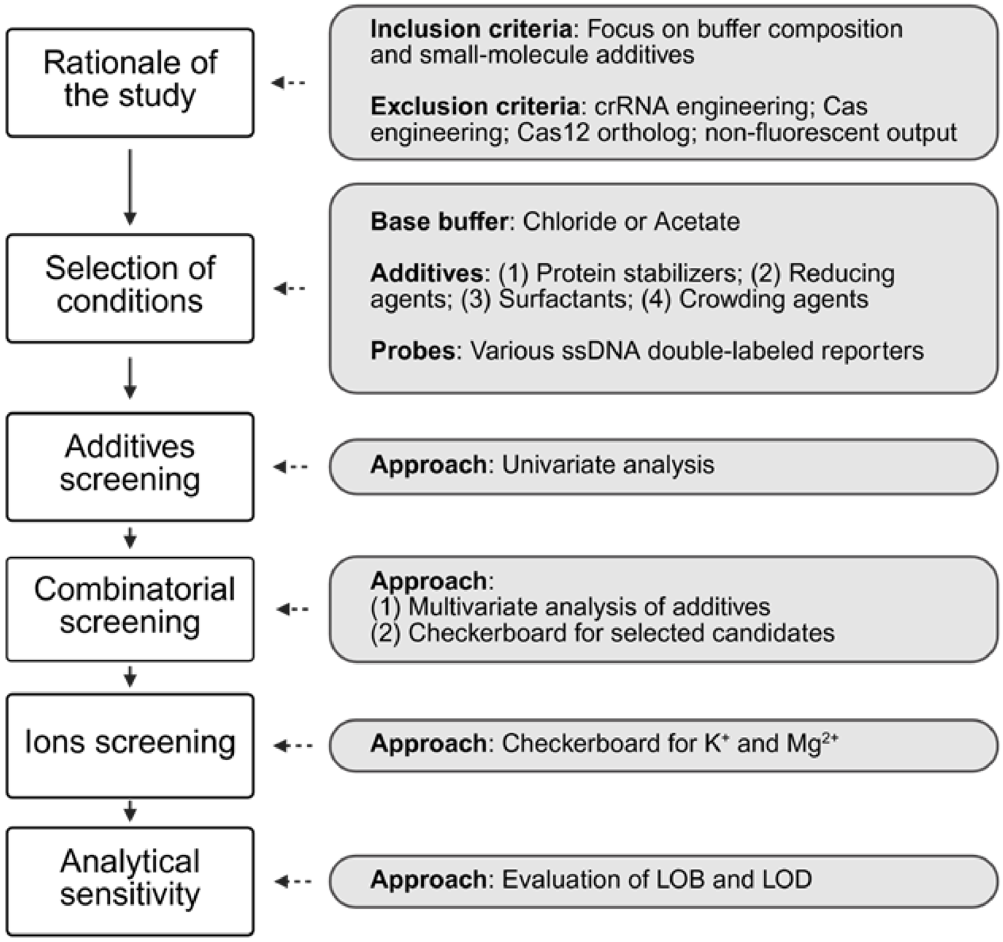


**Figure S1. Schematic workflow for evaluating the effects of buffer composition on LbCas12a trans-cleavage activity.** Relevant studies about Cas12a trans-cleavage activity optimization were retrieved using PubMed, Semantic Scholar, and Research Rabbit. Studies focused on buffer composition and chemical additives were considered, whereas studies involving crRNA engineering, Cas engineering, Cas12 ortholog comparisons, or non-fluorescent outputs were excluded. Two buffer systems (Tris-$Cl^-$ and Tris-$Ac^-$) were selected for downstream evaluation. Reported additives were grouped based on their expected role on enzyme activity, such as protein stabilizers, reducing agents, surfactants, and crowding agents. The experimental workflow followed a sequential and interdependent strategy, progressing from univariate additive screening to combinatorial optimization, ion screening, and analytical sensitivity evaluation, with each stage guided by the results obtained in the preceding step. All assays were performed with the LbCas12a and a specific crRNA against a synthetic double strand oligo DNA (90 bp long). All reactions were performed at least in triplicate, including appropriate non-template, and non-additive controls.

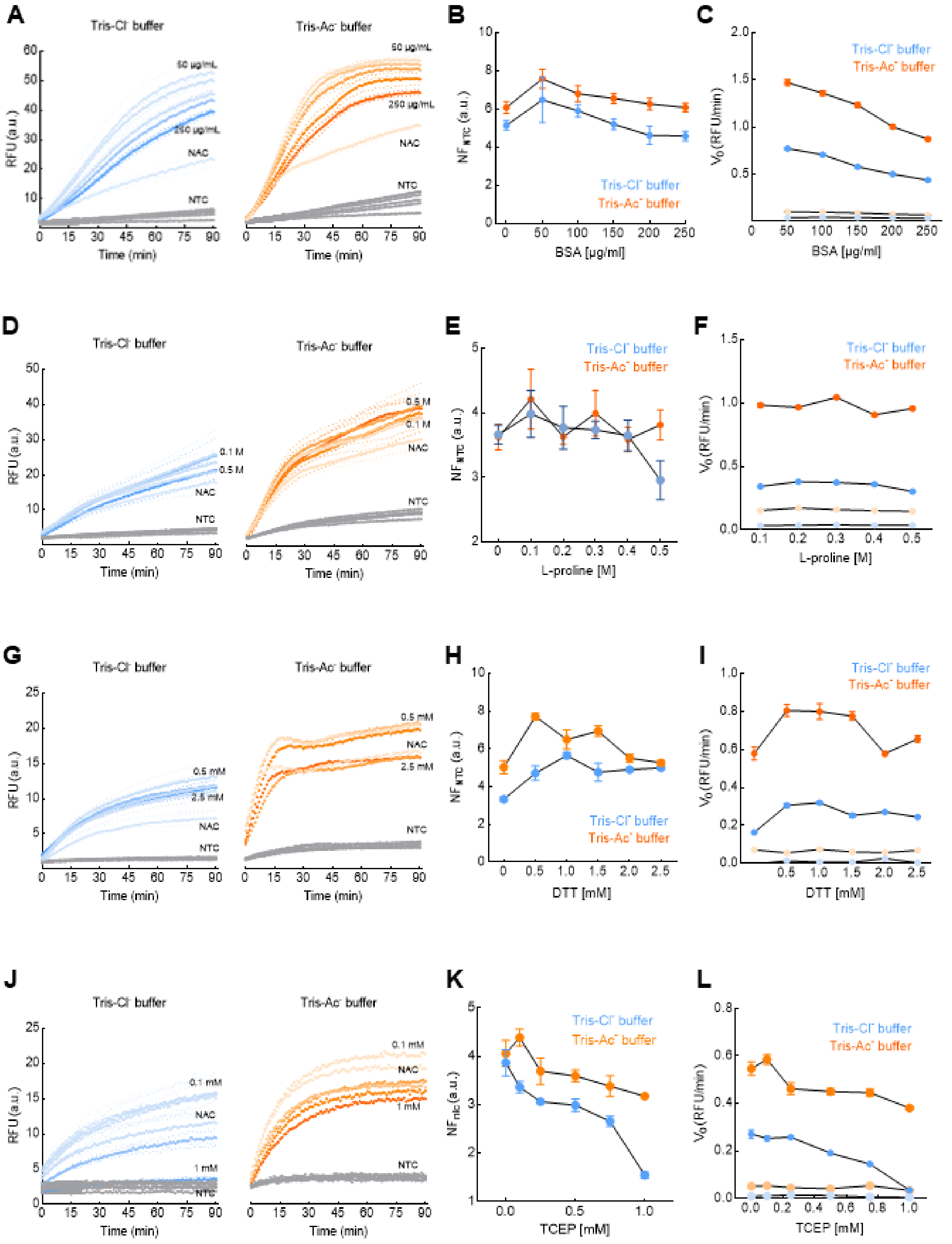


**Figure S2. Univariate screening of different additive groups for the trans-cleavage activity of LbCas12a.** The effect of each additive was evaluated in two different buffer systems: Tris-Cl$^-$ (10 mM Tris-HCl, 50 mM NaCl; blue) and Tris-Ac$^-$ (20 mM Tris-acetate, 50 mM $CH_3CO_2K$; orange). Panels **A–C** correspond to BSA (0 - 250 µg/mL), panels **D–F** to L-proline (0 – 0.5 M), panels **G–I** to DTT (0 – 2.5 mM), and panels **J–L** to TCEP (0 - 1 mM). Real-time fluorescence kinetics are shown in panels **A, D, G,** and **J,** with increasing concentrations of additives compared to reactions without additives and non-template controls (NTC, gray lines). Panels **B, E, H,** and **K** present dose-response analyses of normalized fluorescence ratios ($NF_{ntc}$) relative to the NTC, while panels **C, F, I,** and **L** display the corresponding initial velocity ($V_0$) values derived from the linear phase. Dotted lines represent the standard deviation (n=3). NAC represents the non-additive control. NTC represents the non-template control.

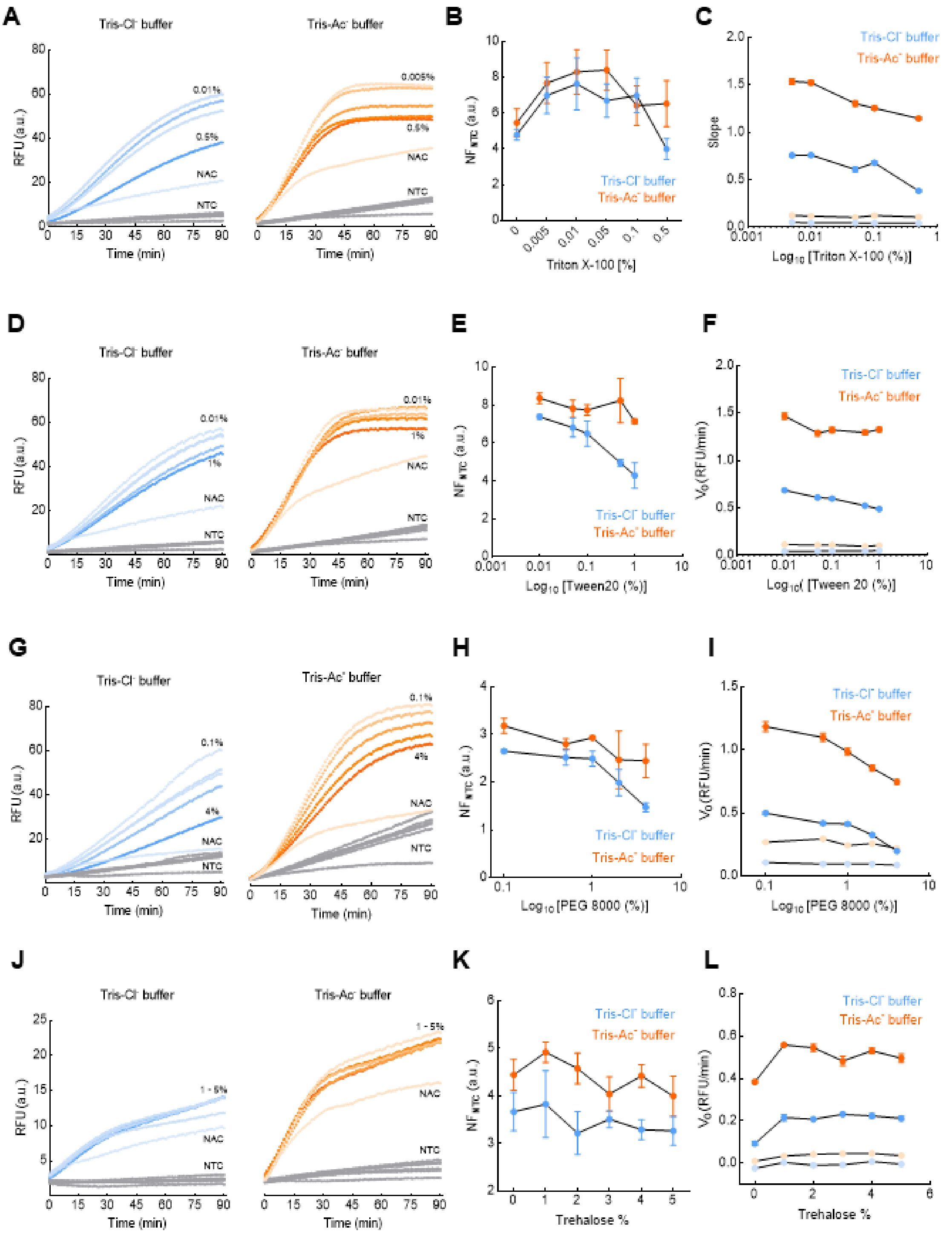


**Figure S3. Univariate screening of different additive groups for the trans-cleavage activity of LbCas12a.** The effect of each additive was evaluated in two different buffer systems: Tris-Cl$^-$ (10 mM Tris-HCl, 50 mM NaCl; blue) and Tris-Ac$^-$ (20 mM Tris-acetate, 50 mM $CH_3CO_2K$; orange). Panels **A–C** correspond to Triton X-100 (0 - 0.5%), panels **D–F** to Tween 20 (0 – 1%), panels **G–I** to PEG 8000 (0 – 4%), and panels **J–L** to Trehalose (0 – 5%). Real-time fluorescence kinetics are shown in panels **A, D, G,** and **J,** with increasing concentrations of additives compared to reactions without additives and non-template controls (NTC, gray lines). Panels **B, E, H,** and **K** present dose-response analyses of normalized fluorescence ratios ($NF_{ntc}$) relative to the NTC, while panels **C, F, I,** and **L** display the corresponding initial velocity ($V_0$) values derived from the linear phase. Dotted lines represent the standard deviation (n=3). NAC represents the non-additive control. NTC represents the non-template control.

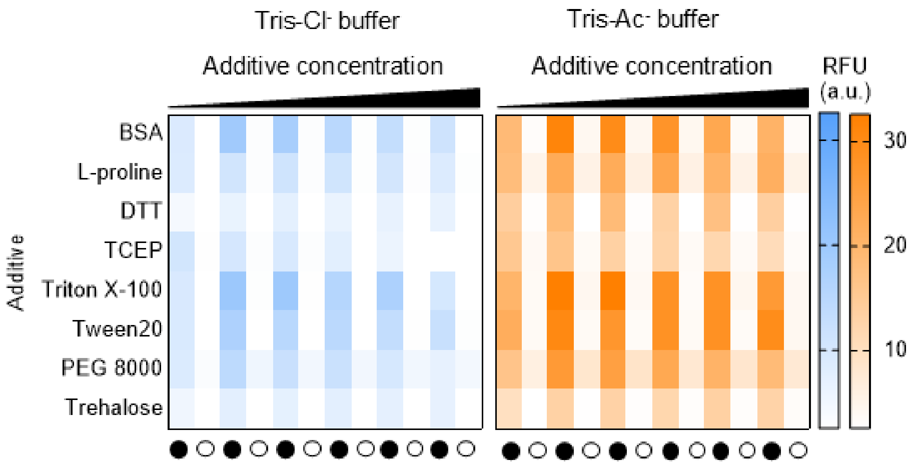


**Figure S4. Summary of fluorescence assessment from the univariate screening.** Heatmap summary of fluorescence values (a.u.) at the 20-min readout time across the tested additives concentration gradients (50 - 250 µg/mL BSA; 0.1 – 0.5 µM L-proline; 0.5 - 2.5 mM DTT; 0.1 - 1 mM TCEP; 0.01 - 1% Tween 20; 0.005 - 0.5% Triton X-100; 0.1 - 4% PEG 8000; and 1 - 5% D-trehalose). For each additive gradient in both buffer systems, non-additive controls (first two columns of each heatmap) were included, and at every concentration both template (black circle) and non-template (white circle) reactions were evaluated.

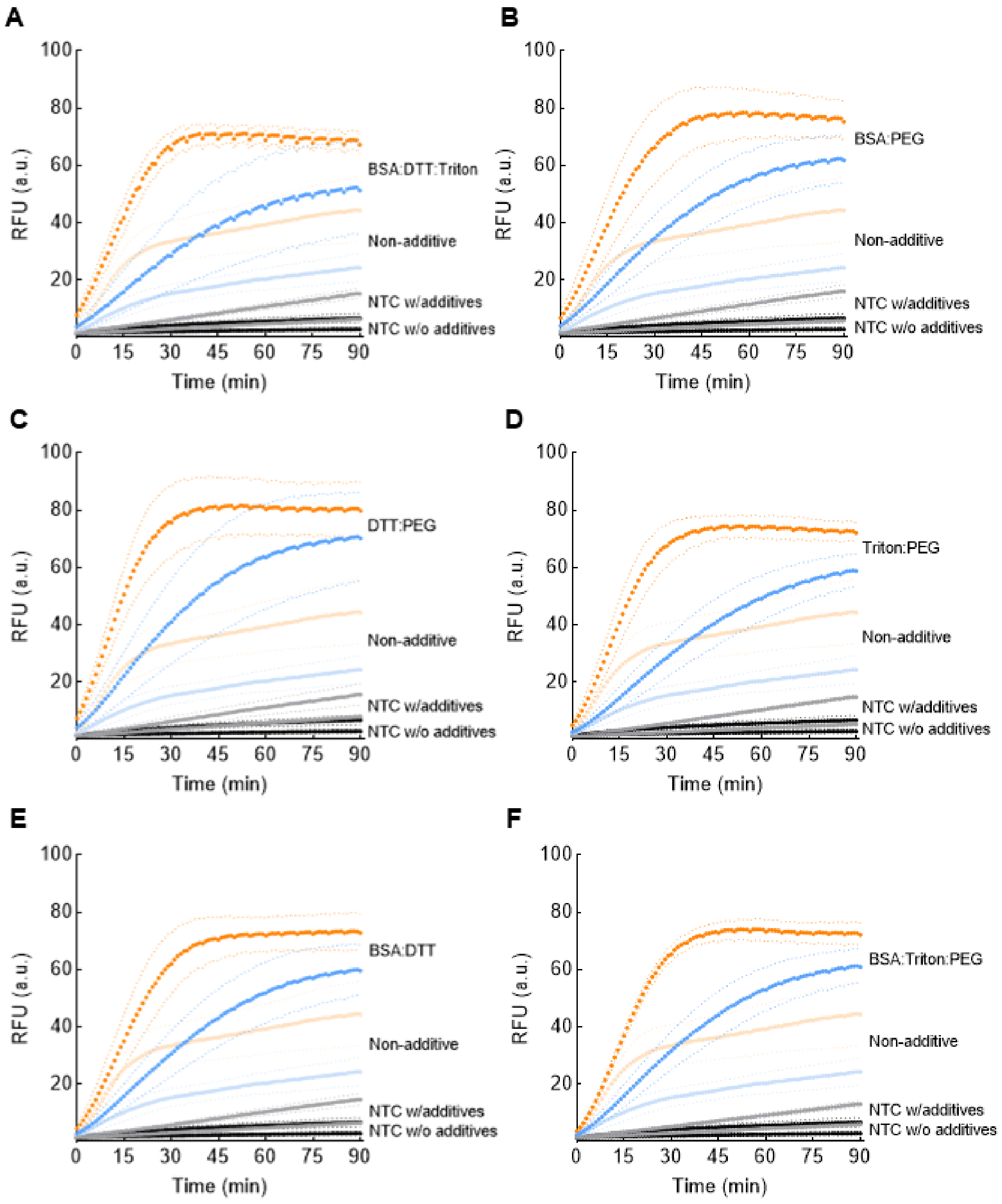


**Figure S5. Combinatorial screening of buffer additives for synergistic enhancement on the LbCas12a trans-cleavage activity.** Real-time fluorescence kinetics for different additives combinations compared to non-additive controls. Tris-Ac$^-$ system yields the highest absolute fluorescence intensity but also induces a considerable time-dependent rise in non-specific background noise (NTC-additive, grey line). Dotted lines represent the standard deviation (n=3).

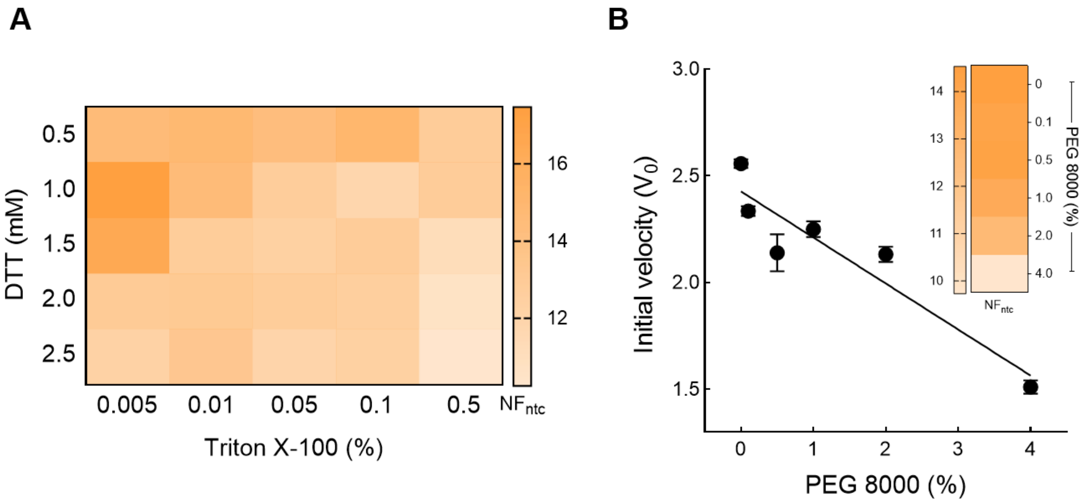


**Figure S6. Combinatorial screening of top additive enhancers: effect of DTT, Triton X-100 and PEG 8000, on the LbCas12a trans-cleavage activity.** **(A)** Checkerboard screening of Triton X-100 and DTT in Tris-$Ac^-$ buffer. Heatmap intensity represents the normalized fluorescence ratio ($NF_{ntc}$) at 20 minutes. A positive trend was observed at low concentrations of both DTT and Triton X-100. **(B)** Dose-response analysis of PEG 8000 on initial velocity ($V_0$) in the Tris-$Ac^-$ buffer supplemented with 1.0 mM DTT and 0.005% Triton X-100. The reaction exhibited an inhibitory effect, with increasing PEG concentrations proportionally reducing $V_0$. The inset heatmap shows $NF_{ntc}$ values obtained across the concentration gradient of PEG.

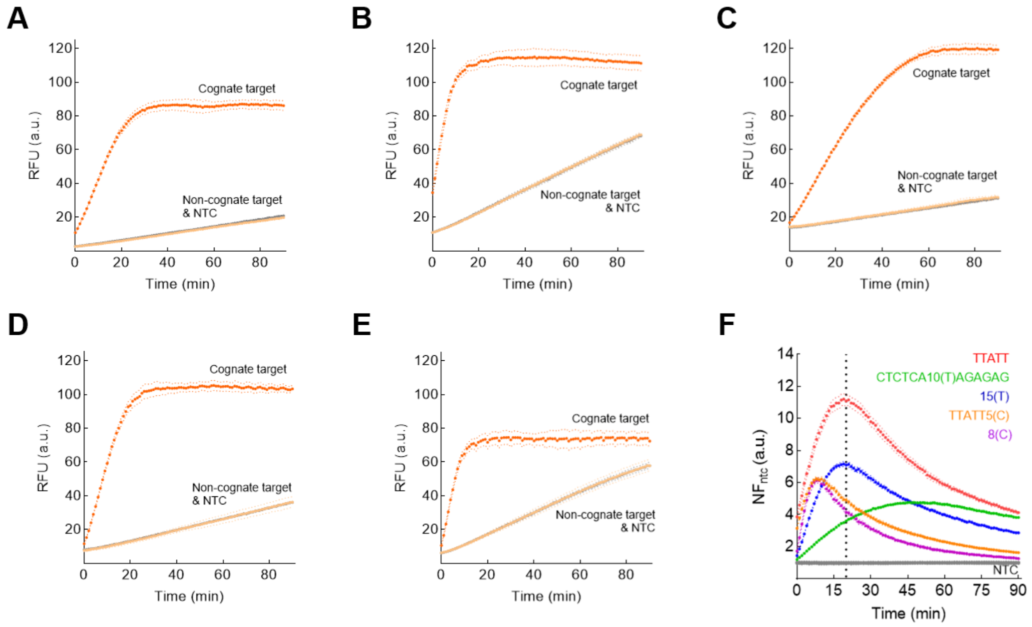


**Figure S7. Effect of reporter probe sequence on the LbCas12a trans-cleavage activity.** CRISPR-Cas reactions were performed in high-salt Tris-Ac-buffer supplemented with DTT and Triton X-100. Panels **A-E** show real-time fluorescence kinetics for cognate and non-cognate targets, as well as non-template control (NTC), using **(A)** the canonical reporter (TTATT), **(B)** the modified canonical reporter (TTATTCCCCC), **(C)** hairpin reporter (CTCTCA-10T-AGAGAG), **(D)** homopolymeric poly-T reporter (15T), **(E)** homopolymeric poly-C reporter (8C). Dotted lines represent the standard error (n=3). **(F)** Same as A-E, but for normalized fluorescence ratios ($NF_{ntc}$). The vertical dotted line (20 min) marks the cutoff of a typical detection window. Dotted lines represent the standard deviation (n=3).

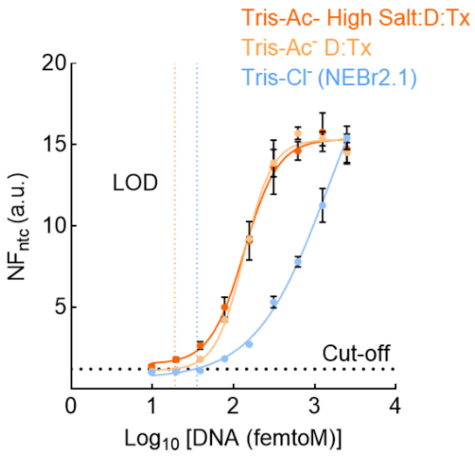


**Figure S8. Limit of detection estimation (LOD).** Normalized fluorescence ratio (NFntc) is presented as a function of target concentration Vertical dotted lines indicate LOD (Limit of Detection) values calculated at 20 minutes of fluorescence measurement, while the horizontal dotted line represents the $NF_{NTC}$ LoB (Limit of Blank, cutoff), estimated at 1.2 a.u.. The LOD for the Tris-Ac- buffer supplemented with DTT, Triton X-100 and high $K^+/Mg^{2+}$ concentrations could not be determined under the present results.

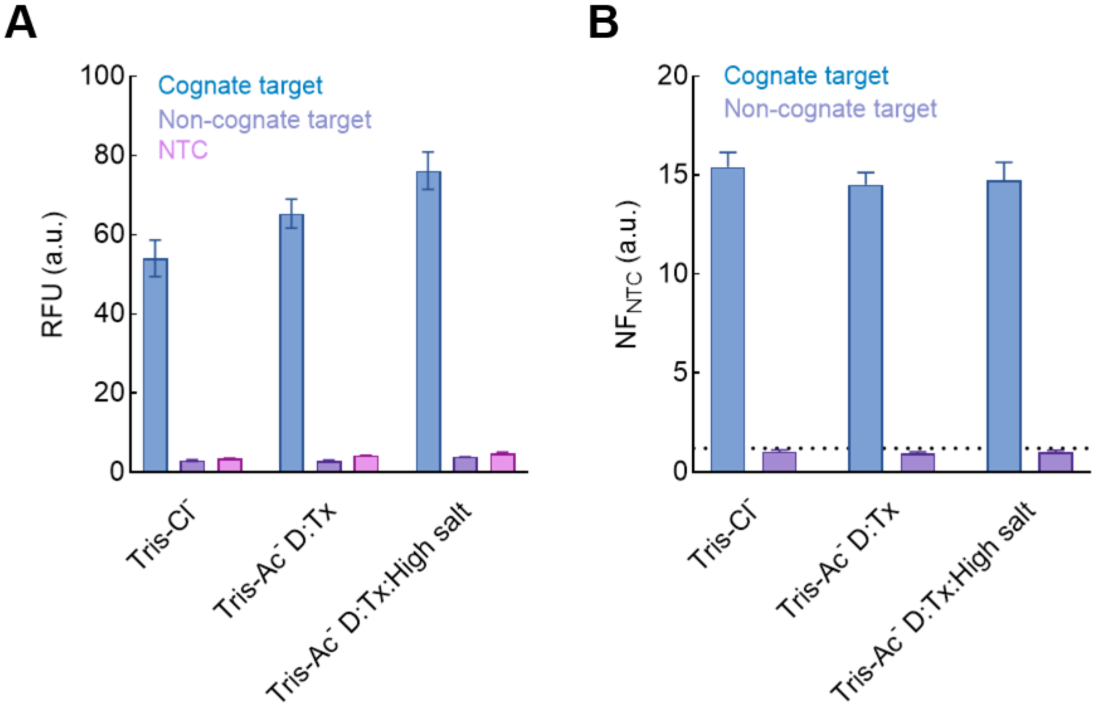


**Figure S9. Specificity of fluorescence signal across buffer conditions.** Fluorescence signal (**(A)** raw RFU and **(B)** normalized $NF_{ntc}$ values) was evaluated across different buffer compositions (Tris-Cl$^-$, Tris-Ac$^-$ DTT:Triton X-100 (D:Tx), and Tris-Ac$^-$ DTT:Triton X-100 with high salt concentration (D:Tx:High salt). In all cases, fluorescent signal was specific to the cognate target, with lower fluorescence observed for non-cognate targets and non-template controls. The dotted line in Panel **B** represents the LOB estimated for all reaction conditions. Data correspond to the highest concentrations tested in the titration gradient for both cognate and non-cognate targets.
